# Horseradish peroxidase as an electrochemical reporter protein for cell-free biosensors

**DOI:** 10.64898/2025.12.17.694927

**Authors:** Aarushi Ruhela, Sahan B. W. Liyanagedera, Nadanai Laohakunakorn, Jamie R. K. Marland

**Affiliations:** School of Engineering, Institute for Integrated Micro and Nano Systems, University of Edinburgh, Edinburgh EH9 3FF, United Kingdom; Centre for Engineering Biology, Institute of Quantitative Biology, Biochemistry and Biotechnology, School of Biological Sciences, University of Edinburgh, Edinburgh EH9 3FF, United Kingdom; Biophoundry Inc., Chapel Hill, North Carolina, United States

**Keywords:** Synthetic biology, Biosensors, Cell-free biosensors, Next-generation biosensors, Electrochemistry, Horseradish peroxidase, Redox enzyme

## Abstract

Cell-free biosensor systems offer a promising platform for portable diagnostics. However, most employ fluorescent reporter proteins that require complex instrumentation and can be affected by photo-bleaching and auto-fluorescence, limiting translatability. Electrochemical reporters do not suffer from these drawbacks. Here, we evaluate horseradish peroxidase (HRP) as a redox enzyme reporter for cell-free biosensor systems. HRP was synthesized in an *E. coli* cell-free transcription-translation system supplemented with hemin, calcium acetate, and commercial disulfide bond enhancers. The electrochemical detection of its activity was established by chronoamperometry, with *H*_2_*O*_2_ as a substrate and tetramethylbenzidine as a redox mediator. Cell-free expressed HRP produced a strong steady state current compared to a catalytically inactive mutant and a no-template control. Kinetic analysis showed a *K*_*m*_ for the cell-free expressed HRP close to that of the native enzyme. To explore the potential of HRP as an electrochemical reporter, we placed it under the control of a tetracycline-responsive regulatory promoter and demonstrated a 3-fold current increase in the presence of anhydrotetracycline. These results support HRP as an electrochemical reporter for cell-free biosensors, offering a practical alternative to optical reporters for future use in handheld analytical devices.

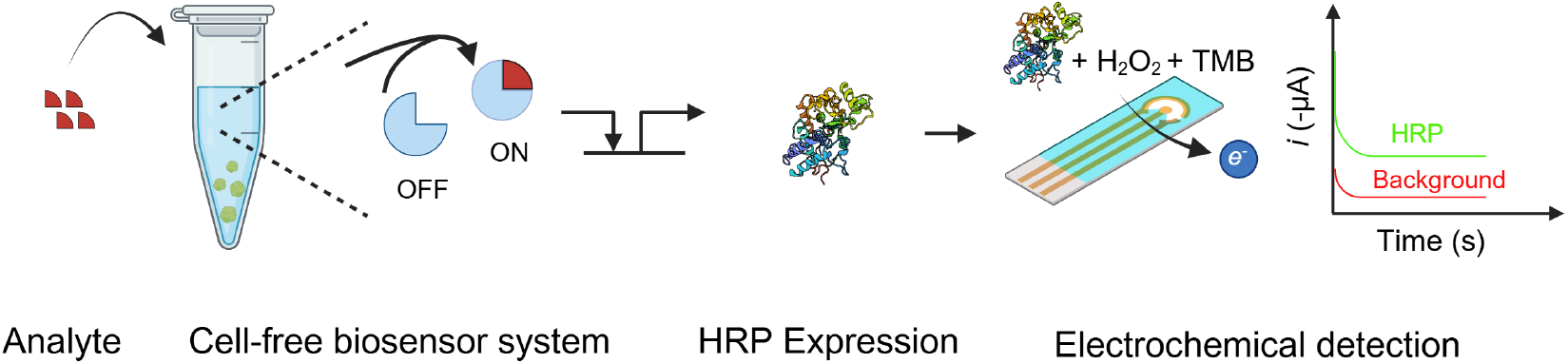

## 1 Introduction

Biosensors are analytical devices that couple a biological recognition element with a physicochemical transducer, providing highly selective, sensitive, low-cost, and rapid readout. These advantages make them attractive for applications ranging from environmental monitoring to point-of-care diagnostics. Advances in synthetic biology have enabled new strategies for biosensor design and construction (*1*). Among these, cell-free protein synthesis (CFPS) systems comprising cellular lysates or reconstituted transcription–translation (TX–TL) machinery enable *in vitro* gene expression without the constraints of cellular viability or regulation (*2, 3*). CFPS platforms can therefore be engineered to detect target analytes and transduce this recognition into a measurable output signal (*4*). Conceptually, cell-free biosensors consist of three functional modules: a detection module that recognizes the analyte, a processing module that transduces the recognition event, and an output module that generates a measurable signal (Figure 1a) (*5*). Traditionally, the output module has relied on fluorescent proteins (e.g., GFP or mCherry) that are commonly used for their visual simplicity, but require bulky optical instrumentation such as fluorimeters and are limited by spectral overlap and background autofluorescence. These reporter systems are primarily suited to laboratory demonstrations and face significant challenges in scalability and real-world deployment.

**Figure 1.**
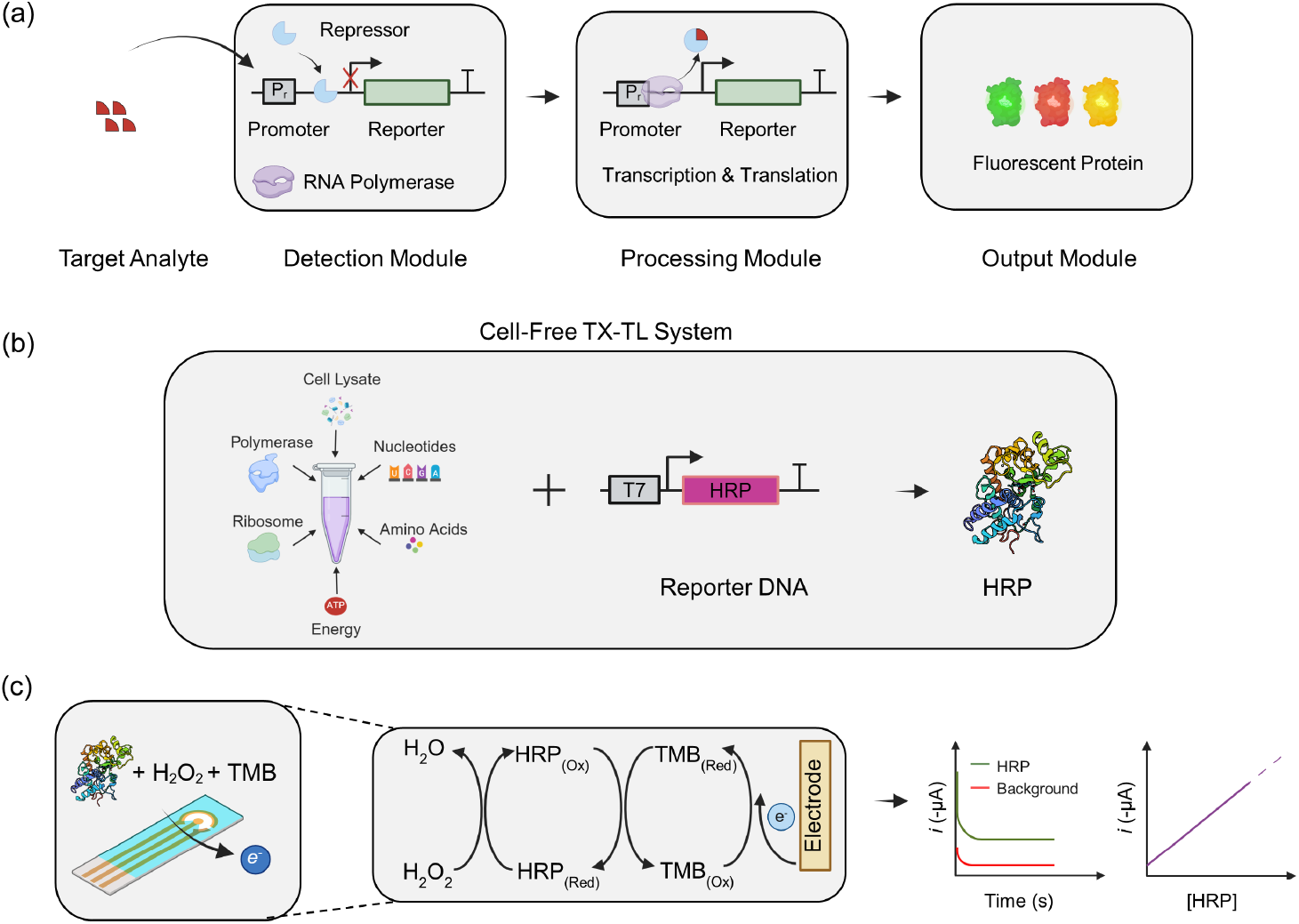
Schematic representation of the cell-free biosensor coupled with electrochemical detection using HRP as a reporter protein. (a) In the cell-free biosensor system, transcription of the reporter gene is repressed in the absence of the target analyte. Upon analyte binding, the repressor is displaced, activating transcription and leading to reporter expression. (b) Schematic representation of HRP expression from reporter DNA templates under a T7 promoter in an *E. coli* cell-free TX–TL system. (c) The electrochemical detection mechanism relies on HRP-catalysed redox cycling of TMB in the presence of H_2_O_2_, producing a measurable current at the electrode surface. The output signal is recorded as a reduction current (*i*) over time, which increases proportionally with HRP concentration, enabling quantitative detection.

Electrochemical readouts address these issues by offering several advantages, particularly for medical diagnostics and integration with digital health platforms (*6*). They are inherently quantitative, can be miniaturised for point-of-care applications, and are compatible with portable, low-power electronics. Electrochemical detection has been used extensively in the past to detect cells, proteins, nucleic acids, and small molecules (*7*) and, in some instances, redox enzymes have been employed to establish the interface between the biological events and electronic signals (*8, 9*).

In addition, enzyme-based reporters provide a substantial amplification advantage over fluorescent proteins due to their high substrate turnover rate, leading to faster readouts and a lower limit of detection (LoD) (*10*). Enzymatic reporters such as,*6*-galactosidase and luciferase, which have been used in cell-based systems, generate colourimetric or luminescent outputs but are typically limited to laboratory utility or only give qualitative results (*11, 12*). Therefore, in this work, we explored Horseradish peroxidase (HRP) as a redox enzyme reporter for transducing the input of a cell-free biosensor into quantifiable electronic output signals.

Native HRP is a glycosylated heme-containing monomeric protein of approximately 40 kDa in size, containing four disulfide bonds (*13*). It has a well-established reagent chemistry with a broad range of readily available analytical tools and is widely used in biosensing and diagnostic applications for immunohistochemistry, enzyme-linked immunosorbent assay, in-situ hybridisation, DNA biosensors and blot analysis (*14*). Studies have shown that the recombinant expression of active HRP in *E. coli* is possible but presents several challenges. It typically requires refolding steps or co-expression of folding chaperones and proper disulfide bond formation to restore enzymatic functionality (*15–18*), and the heme co-factor must be efficiently supplemented or biosynthesised for proper catalytic assembly (*19*). Despite these challenges, active HRP can be expressed in CFPS by providing the necessary folding conditions and co-factors (*20, 21*).

In this work, we translated HRP in an *E. coli*-based cell-free TX-TL system (Figure 1b). We further established an amperometric detection for this cell-free expressed HRP using screen-printed gold electrodes (Figure 1c). The expressed HRP catalyses the reduction of hydrogen peroxide (H_2_O_2_) to water, and oxidises the mediator substrate, 3,3^*‘*^,5,5^*‘*^tetramethylbenzidine (TMB), producing an electrochemically active product. The measured current output is generated from the reduction of HRP-oxidised TMB at the electrode surface, which increases proportionally with increasing HRP concentration. The resulting chronoamperometry response provides a quantitative electrochemical readout of analyte-dependent transcriptional activity (Figure 1d). To validate the utility of using HRP as a cell-free biosensor reporter, we demonstrated the expression of HRP under the control of a tetracycline-responsive regulatory promoter for anhydrotetracycline (aTc) sensing.

## 2 Results

### 2.1 Amperometric detection of HRP

We first established an electrochemical protocol for HRP detection based on the HRP-mediated generation of oxidised TMB (TMB_*ox*_), which is subsequently reduced at the electrode. Purified native HRP was used in these initial experiments to validate the protocol before applying it to the HRP expressed from the cell-free system. Cyclic voltammetry (CV) of the substrate solution containing H_2_O_2_ and TMB was performed to characterize the system. The anodic sweep was limited to +0.1 V to avoid direct electrochemical oxidation of TMB at the electrode. Substrate alone showed a negligible Faradaic reduction current, consistent with the absence of TMB_*ox*_ (Figure 2a). In contrast, incubation of the substrate with HRP (5 mUnits/mL) for 2 minutes produced a pronounced reduction current, with a cathodic peak at approximately +0.06 V, corresponding to the reduction of enzymatically oxidised TMB (Figure 2a). The highest signal-to-background separation occurred at 0.0 V, which was therefore selected for chronoamperometry (CA) to quantify HRP activity. CA measurements showed an initial transient current, followed by the development of a steady-state reduction current. The steady-state current of substrate-only control was *-*8.3 *±* 1.1 nA, whereas the presence of HRP (5 mUnits/mL) increased the current to *-*302.6 *±* 25.9 nA, confirming that the CA signal reflected HRP activity (Figure 2b).

**Figure 2.**
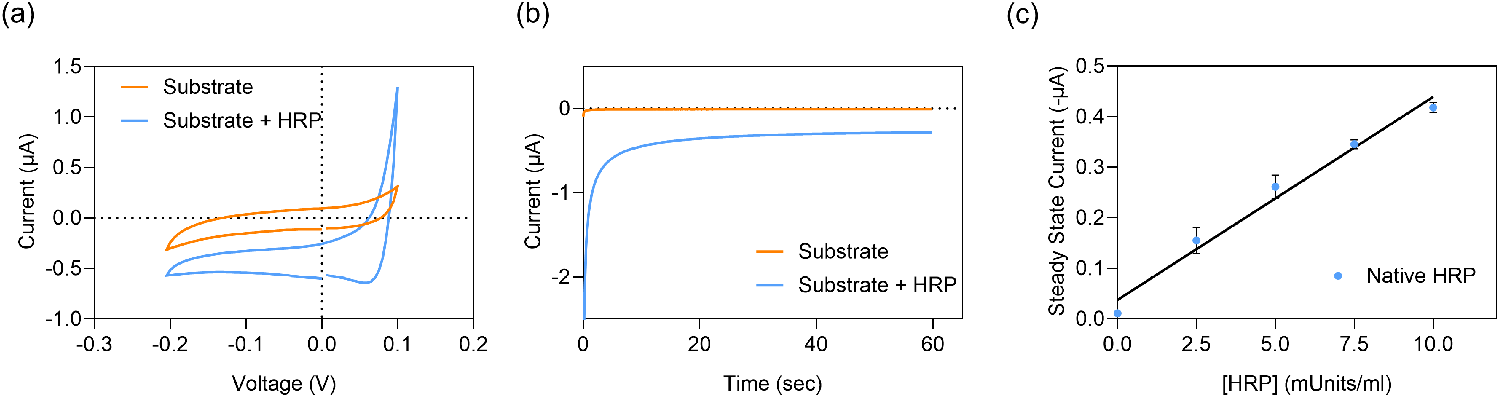
(a) Cyclic voltammetry (CV) of substrate solution without and with HRP (5 mUnits/mL), recorded over a potential range of 0.2 to +0.1 V. (b) Chronoamperometry (CA) of substrate solution without and with HRP (5 mUnits/mL), performed at an applied potential of 0.0 V for 60 s (c) Calibration curve of steady-state CA current as a function of HRP concentration, showing a linear response (*R*^2^ = 0.95). Calibration was performed under the same CA conditions. All measurements were taken after a 2-minute incubation of HRP with substrate prior to electrochemical readings. Error bars represent the standard error of the mean and replicates from n = 3 experiments.

Then, to determine the sensitivity and LoD of the assay, steady-state CA currents were measured at varying concentrations of HRP. As shown in Figure 2c, a clear linear relationship between HRP concentration and steady-state current was observed, enabling quantitative sensing. The sensitivity to HRP was 40.17 *±* 2.43 nA per (mUnit/ml) with an offset of 0.04 *±* 0.01 *µ*A. The LoD (three times the standard deviation of the blank signal, divided by sensitivity) was 0.14 mUnits/mL.

### 2.2 Cell-free expression and electrochemical detection of HRP

Expression of HRP in the cell-free TX-TL system was first optimised by varying the incubation temperature, hemin, calcium concentration, and screened for the effects of commercial disulfide bond enhancers (Supplementary Fig S2, S3, S4). The cell-free expression and functional validation of HRP were initially assessed using an optical assay with H_2_O_2_ and TMB as a chromogenic substrate. In the presence of HRP, TMB is converted into a coloured product with absorbance at 450 nm when quenched with acid. A significant increase in absorbance was observed in the wild-type (WT) HRP expressing reaction (WT), whereas negligible signals were detected in the no-DNA template control (NTC) or a catalytically inactive mutant (H170A) (*22*) (Figure 3a). Absorbance readings for HRP (3.014 *±* 0.074) were significantly greater than those of NTC (0.064 *±* 0.001) or H170A (0.066 *±* 0.002). After confirming HRP expression using the optical method, we next evaluated the electrochemical detection of cell-free expressed HRP. The steady-state reduction current was markedly higher in the HRP sample (WT) (52.56 *±* 1.58 nA) compared to both the NTC (0.73 *±* 0.13 nA) and the H170 mutant (0.55 *±* 0.06 nA), confirming that the observed signal originated from the enzymatic activity of HRP (Figure 3b). No significant difference was observed between NTC and H170A in both optical and electrochemical methods, indicating that the signal was specific to the functional peroxidase and not due to background activity from the cell-free extract, and that the H170A variant lacks measurable enzymatic activity. SDS-PAGE showed successful translation of both H170 and WT HRP (Supplementary Fig. S1). Furthermore, the electrochemical results were consistent with those obtained from the optical assay, demonstrating the reliability of the electrochemical platform in detecting cell-free expressed HRP.

**Figure 3.**
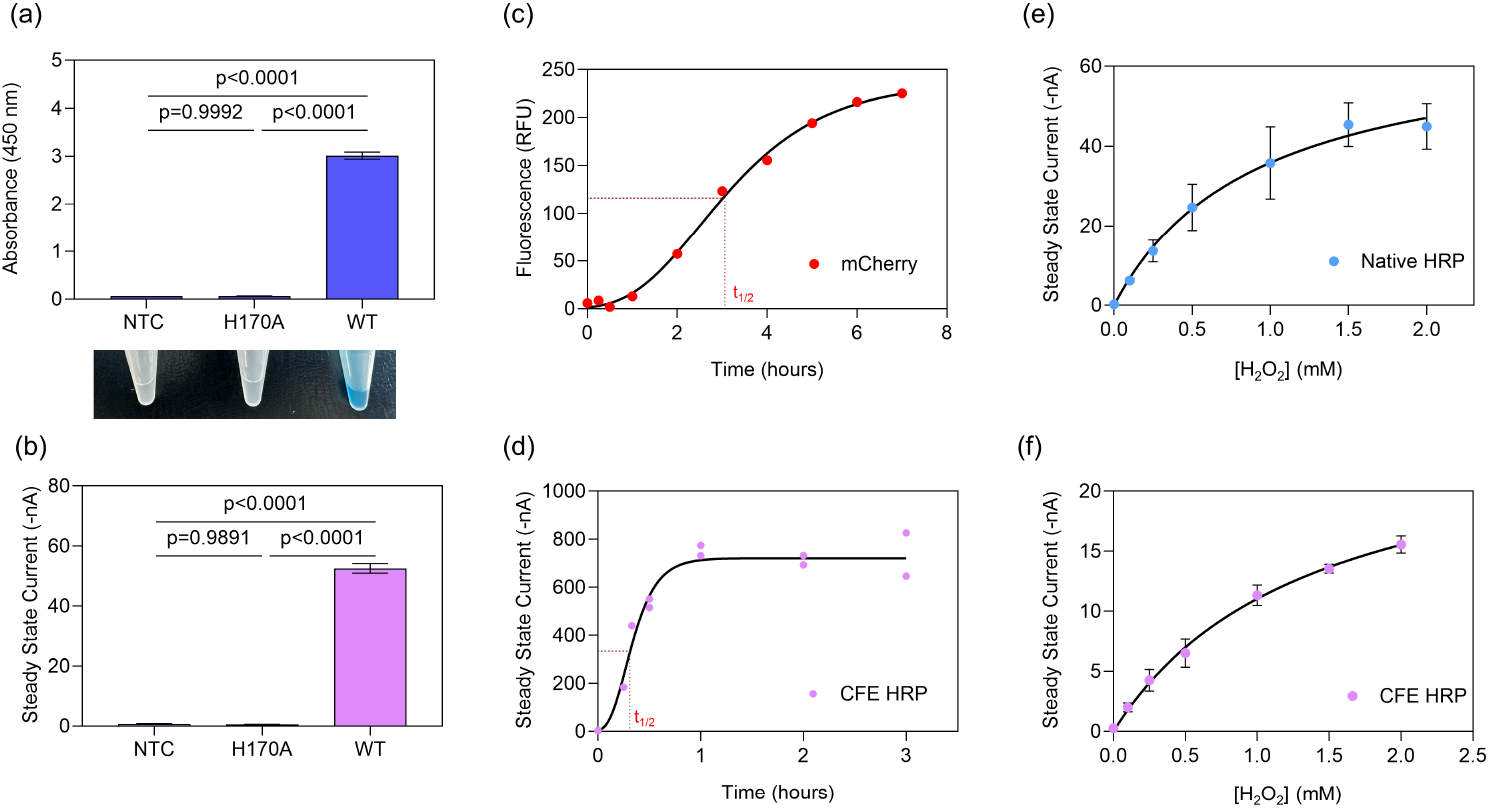
Cell-free reactions were assembled with 10 nM DNA template encoding wild-type HRP (WT) or the catalytically inactive mutant H170A; no-DNA-template reactions (NTC) served as negative controls. (a) Spectrophotometric detection of HRP activity using TMB substrate for NTC, H170A, and WT, with photographs of the corresponding reaction tubes shown below. (b) Electrochemical detection using chronoamperometric steady-state current for NTC, H170A, and WT reactions. (c) Time-course of mCherry fluorescence used as a TX-TL reporter, with the half-maximum time (*t*_1*/*2_) indicated. (d) Time-course of steady-state chronoamperometric current for WT HRP expression in the cell-free system, with *t*_1*/*2_ indicated. Michaelis–Menten analysis of (e) purified native HRP (4 mUnits/mL) and (f) cell-free expressed (CFE) HRP (10 *µ*L of 10*⇥* diluted reaction product), determined from steady-state chronoamperometric currents at varying H_2_O_2_ concentrations and fixed TMB (260 *µ*M). Each data point represents the mean of *n* = 3 technical replicates.

Previous work has shown that the enzymatic reporters can mature more rapidly than fluorescent proteins in the cell-free systems (*10*). To explore this, we measured the expression kinetics of HRP and mCherry as a fluorescent protein and fitted the data using an asymmetric sigmoidal growth model (Gompertz) that captures both the initial lag phase and the eventual saturation. The time to reach half the maximum signal was estimated to be 19 minutes for HRP, compared with 184 minutes for mCherry (Figure 3c,d). This result is consistent with the previous reports of enzymatic reporters providing substantially faster readout than fluorescent proteins, and is advantageous for faster signal generation in biosensing applications.

Finally, to assess the functional quality of cell-free expressed HRP, we performed a Michaelis–Menten kinetic analysis using native HRP and cell-free expressed HRP (Figure 3e). The steady-state currents were recorded at varying H_2_O_2_ concentrations with a fixed concentration of TMB. For native HRP, the resulting saturation curve showed characteristic saturating behaviour, with an apparent *K*_*m*_ of 0.91 *±* 0.96 mM. The cell-free expressed HRP displayed a comparable kinetic profile, yielding a *K*_*m*_ of 1.37 *±* 0.48 mM (Figure 3f).

### 2.3 Electrochemical detection of HRP as a reporter protein

Finally, to demonstrate that HRP can function as a reporter protein from a cell-free biosensor system, its expression was placed under the control of a regulated promoter. A commonly used tetracycline-inducible system was selected to validate whether modulation of gene expression can be reliably transduced into a measurable electrochemical signal (*23–25*). The tet-operator (tetO) sequence was positioned downstream of a T7 promoter driving the expression of HRP. Recombinant TetR protein that recognizes and binds the tetO site was added to the cell-free reaction, and it represses basal HRP transcription. Binding of the inducer aTc to TetR releases it from the tetO site, thereby permitting HRP expression (Figure 4a). Cell-free reactions were prepared with the addition of TetR and/or aTc. Initially, to optimize the template design, mCherry expression was regulated with this promoter (Supplementary Fig S5). It was then implemented with HRP as a reporter. As shown in Figure 4b, CA measurements revealed a clear difference in signal between the repressed (OFF) and induced (ON) states. A small reduction current of *-*8.31 *±* 0.45 *µ*A was observed in the absence of aTc (TetR-mediated repression), while a 3-fold higher current of *-*21.06 *±* 0.16 *µ*A was measured upon induction, indicating active HRP expression (Figure 4b). A similar pattern was observed from the colourimetric readout (Supplementary Figure S6). Increasing aTc concentrations led to a dose-dependent induction of HRP expression (Supplementary Figure S7). Together, these results confirm that the synthetic regulatory promoter functioned as intended and that HRP expression was regulated by the tetO system. The successful chronoamperometric detection of HRP activity demonstrates that HRP functions effectively as an electrochemical reporter in the cell-free TX-TL systems, providing a direct and quantifiable connection between the sensing input and the electrical output.

**Figure 4.**
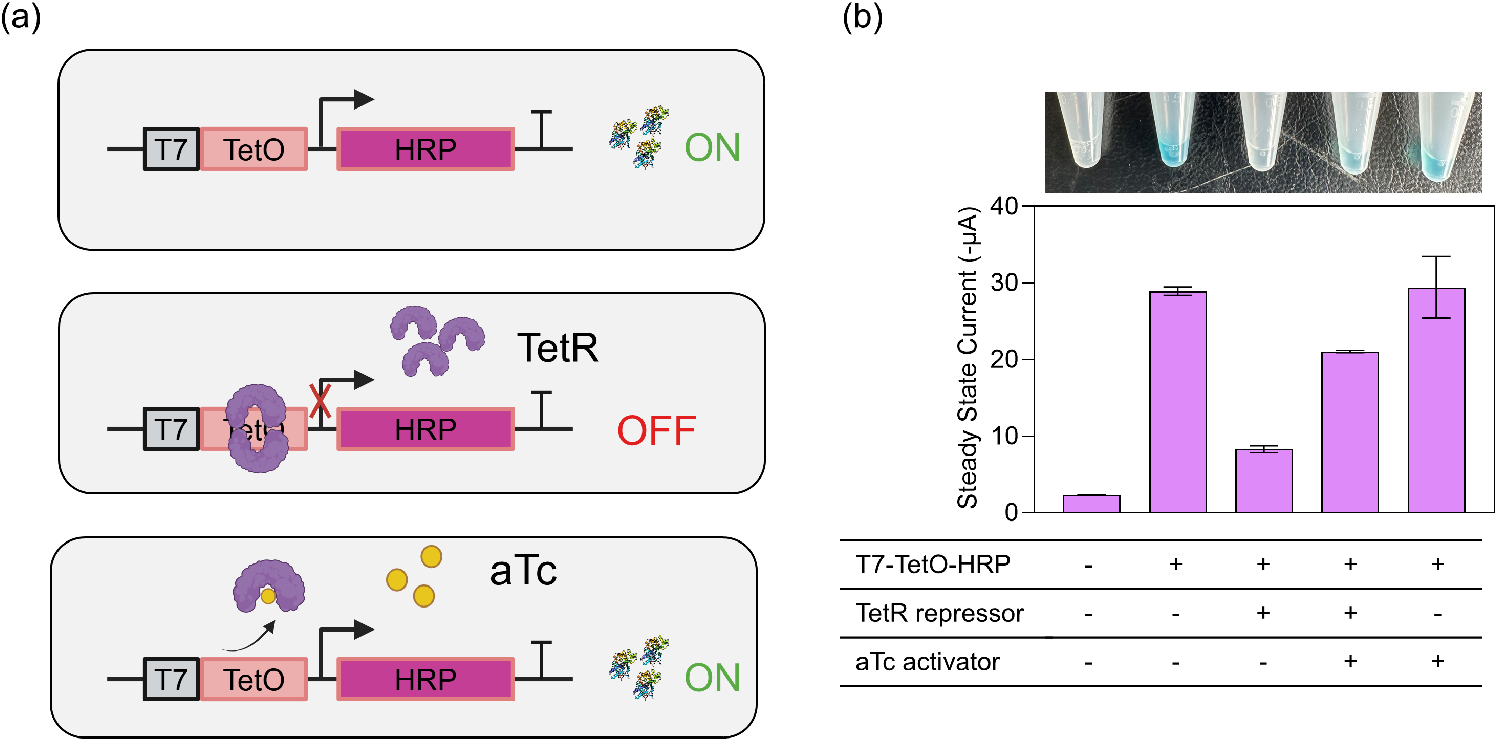
(a) Schematic of the TetR-regulated HRP reporter system. In the absence of TetR, HRP is expressed (ON state). The binding of TetR to the TetO operator represses transcription (OFF state). The addition of aTc inhibits TetR, restoring HRP expression (ON state). (b) Electrochemical detection of HRP activity in the cell-free system, where reactions were set up with 10 nM DNA and/or 100 nM TetR and/or 1 *µ*M aTc. A photograph of colourimetric TMB oxidation (top) and the bar graph of steady-state CA currents (middle) are shown.

## 3 Discussion

The goal of this work was to combine the cell-free TX-TL biosensor system with a fast, sensitive and selective electrochemical response using HRP as a reporter protein that can solve the issues of photo-bleaching and auto-fluorescence associated with conventional fluorescent reporters. This can increase the utility of the cell-free biosensor system for deployment in practical applications.

In this study, HRP was successfully expressed in a CFPS system, consistent with previous work by Zhu *et al*. and Park *et al*., who demonstrated the functional expression of HRP in a cell-free platform (*20, 21*). Similar optimal expression conditions of 20 °C temperature, 15 *µ*M hemin were observed in our work, whereas the optimal Ca^2+^ concentration was found to be lower at 0.5 mM when optimized alongside Mg^2+^ (Supplementary Figure S1, S2, S3). Importantly, the *K*_*m*_ observed from cell-free expressed HRP (1.37 *±* 0.48 mM) was similar to the native HRP (0.91 *±* 0.96 mM), and is also consistent with reported literature values of 0.7 *±* 0.2 mM (*26*). The close agreement between these values confirms that the cell-free product is catalytically active and maintains substrate affinity similar to the purified enzyme. Even in the absence of native glycosylation, which under native conditions supports folding, solubility, and stability (*27*), the cell-free expressed HRP shows close agreement with the purified HRP kinetics despite lacking these modifications in the *E. coli* cell-free system. These findings advance the utility of HRP as a reporter enzyme for cell-free biosensing applications.

CFPS platforms offer various advantages over conventional methods used for biosensing. However, they also have limitations. The stability of cell-free reactions outside laboratory settings has been a challenge and can be mitigated by employing freeze-dried extracts, as demonstrated by Pardee *et al*. (*23*). Furthermore, most CFPS biosensor platforms are laboratory-based because of the dependency on bulky equipment for fluorescent reporter detection. Certain portable formats based on smartphone cameras are available; however, their performance is likely constrained by variability in image acquisition and by requirements for optical filtering and operation in darkness (*28*). In many other instances, cell-free biosensors are applied for on-field testing using paper-based detection, which is straightforward but does not support quantitative readout (*29*). However, electrochemical sensing has the potential to solve both these issues by utilising portable electrodes and a compact potentiostat, comparable to those used in personal glucose monitors, giving a quantitative readout which can be readily integrated into the point-of-care diagnostics platform.

Electrochemical output modules from cell-free TXTL biosensor systems are relatively underexplored. However, in addition to this work, some other approaches have been demonstrated previously. One such example is the use of a nucleic acid-based output module, which relies on synthetic DNA tagged with a redox label such as methylene blue or ferrocene, detected using pulsed amperometry. Tagged single-stranded DNA may be immobilized at an electrode, enabling detection of RNA transcribed within the system and bound to the labelled probe (*30*). Alternatively, untagged single-stranded DNA may be immobilised at the electrode and can capture tagged DNA following its release by translated restriction enzymes (*24*). While these readout methods offer high specificity, they require costly tagged DNA and lack the amplification provided by enzyme-based output modules. Another approach uses an enzyme-based module, in which the cell-free system expresses a reporter enzyme, such as invertase, trehalase, and *β*-galactosidase, that generates glucose as a product (*31–34*). The glucose is then measured using an electrochemical personal glucose meter. This approach enables enzymatic signal amplification using well-established and inexpensive readout technology, but it is vulnerable to distorted measurements due to reaction mixture metabolic activity (such as glycolysis) and existing glucose within test samples. Use of HRP as an electrochemical enzymatic reporter may avoid these potential sources of interference and allow direct redox coupling between the expressed reporter and the electrode. Finally, *β*-galactosidase has also been adapted for electrochemical detection by coupling its enzymatic activity to other electroactive products, which can be monitored through redox-based current changes, enabling the use of *β*-galactosidase in a miniaturized and electronic biosensing format (*35*).

Looking forward, the work presented is well-positioned for integration with a broad range of biosensing applications (*36*). Coupling of HRP as a reporter protein to existing or new biosensing modules would help the transition to electrochemical readout, which may facilitate the implementation in field-deployable settings (*37*). There is also scope in exploring alternative redox reporters or more stable substrates other than TMB to broaden the toolkit available for cell-free biosensors, offering new avenues for optimising both sensitivity and operational stability (*26*).

## 4 Conclusion

The HRP-based electrochemical reporter system described here combines enzymatic amplification with the portability and simplicity of electrochemical detection, offering a compelling alternative to traditional optical reporters in both research and applied contexts. We believe that this work will open new opportunities to expand the use of cell-free systems for on-site biosensing and diagnostics without the need to rely on bulky optical equipment. With the increasing integration of bioelectronics, this study also bridges the gap between synthetic biology and electronics, providing a foundation for the future development of lab-on-chip electronic devices.

## 5 Methods

### 5.1 DNA preparation

Synthetic gene fragments for the WT HRP, H170A mutant, T7tetOHRP and T7tetOmCherry were synthesized by Integrated DNA Technologies (IDT) (sequences are reported in Supplementary Sequence Information) and assembled into the T7p14 expression vector using the Gibson Assembly® Master Mix (New England Biolabs (NEB)), following the manufacturer’s instructions. The resulting plasmid constructs were sequence-verified by Sanger sequencing (Genewiz) to confirm correct assembly. Subsequently, plasmid DNA was prepared using the ZymoPURE Plasmid Maxiprep Kit (Zymo Research) and linear DNA was PCR amplified using primers that bind 250 base-pairs upstream and downstream (Supplementary Table S1) of the inserts using Phusion™ High-Fidelity DNA Polymerase (NEB) following the manufacturer’s instructions. All the DNA samples were further purified using a DNA Clean & Concentrator-5 Kit (Zymo Research) to remove residual salts and enzyme contaminants before downstream applications.

### 5.2 Crude extract preparation

Cell extracts were prepared by growing *E*.*coli* BL21 cells at 37 °C to an OD_600_ of 1.8. The protocol for the downstream processing from Liyanagedera *et al*. was followed (*38*).

### 5.3 Cell-free TX-TL expression

Cell-free TX-TL reactions of 10 *µ*L volume were assembled with 10 nM plasmid or linear DNA and supplemented with appropriate concentrations of additives including NTPs, amino acids, tRNAs, salts, and crowding agents (Supplementary Table S2). In addition to these, the reaction mixtures for HRP were further supplemented with 15 *µ*M hemin, 0.5 mM calcium acetate, and a commercial disulfide bond enhancer kit (PURExpress® Disulfide Bond Enhancer, NEB) to facilitate proper folding and catalytic activity of the HRP enzyme. The reactions were incubated at 20°C for 8 hours for HRP and for mCherry at 29°C for 12 hours. The reactions for the tet-regulated promoter were supplemented with 100 nM of recombinant TetR protein (HY-P71520, MedChem) and/or 1 *µ*M of anhydrotetracycline hydrochloride (94664, Merck).

### 5.4 Optical detection of HRP activity

The activity of the expressed HRP was confirmed using a conventional colourimetric assay based on the oxidation of TMB in the presence of hydrogen peroxide. Briefly, 90 *µ*L of commercial TMB substrate solution (T4444, Merck) was mixed with 10 *µ*L of diluted cell-free reaction product and incubated for 10 minutes at room temperature, followed by quenching with 50 *µ*L of 1 M H_2_SO_4_. Absorbance was measured at 450 nm using a FLUOstar Omega plate reader (BMG Labtech). Background absorbance from substrate-only controls was subtracted from sample readings prior to data analysis.

### 5.5 Electrochemical detection of HRP activity

Electrochemical measurements were performed using an EmStat3 Blue or Palmsens4 potentiostat (PalmSens BV) with commercial screen-printed electrodes (DRP-C223BT-U75, DropSens) consisting of a 1.6 mm diameter of gold working electrode (WE), silver reference electrode (RE), and gold counter electrode (CE). All potentials are reported relative to the Ag pseudo-reference electrode. Electrochemical cleaning of the electrode surface was performed using 0.1 M H_2_SO_4_ with CV carried out at a scan rate of 0.1 V/s with a step potential of 0.002 V over a range of 0.0-1.6 V before taking any further readings. Later, CV was carried with 40 *µ*L of samples of interest at a scan rate of 0.1 V/s with a step potential of 0.005 V, over a range of -0.2 to +0.1 V. TMB substrate (T4444 or T3405, Merck) was supplemented with 10 mM KCl, and native HRP (P8375, Merck) was used for these initial experiments. CA was performed at an applied potential of 0.0 V (vs. Ag) for 60 s with a sampling interval of 0.1 s, and steady-state current (averaged current readings from 50-60 s) was evaluated.

### 5.6 Fluorescence detection of cell-free reporters

For mCherry expression, 10 *µ*L reactions were set up in a black 384-well plate and incubated at 29°C for 12 hours in a Biotek Synergy H1 plate reader, and the signal was read every 2 minutes with orbital shaking using excitation of 579 nm and emission at 616 nm with a gain setting of 50.

### 5.7 Statistical analysis

Data were analysed using GraphPad Prism version 10.6.1. Error bars represent the standard error of the mean (SEM) from technical replicates (*n* = 3), unless otherwise stated. Statistical significance was assessed using one-way ANOVA with post hoc Tukey’s correction. Non-linear regression analyses were performed using the Michaelis–Menten model for enzyme kinetics and the Gompertz model for TX–TL kinetics of HRP and mCherry.

## Supporting information

Supplementary Sequence Information

## 6 Author Contributions

A.R., S.B.W.L., N.L. and J.R.K.M. designed the experiments. A.R. performed the experiments and analysed the data. S.B.W.L. assisted with the preparation of the cell-free extract. N.L. and J.R.K.M. supervised the project. All authors contributed to writing and editing the manuscript.

## Acknowledgement

A.R. is supported by a PhD studentship from University of Edinburgh. N.L., S.B.W.L. are supported by a UKRI Future Leaders Fellowship to N.L. (MR/V027107/1). The authors gratefully acknowledge technical support from the School of Engineering and School of Biological Sciences and the Centre for Engineering Biology at the University of Edinburgh.

