## Supplementary Information for "Horseradish peroxidase as an electrochemical reporter protein for cell-free biosensors"

### Supplementary Information
### Horseradish Peroxidase as an electrochemical reporter protein for cell-free biosensor systems

Aarushi Ruhela, Sahan B. W. Liyanagedera, Nadanai Laohakunakorn, Jamie R. K. Marland

#### 1 Supplementary Figures

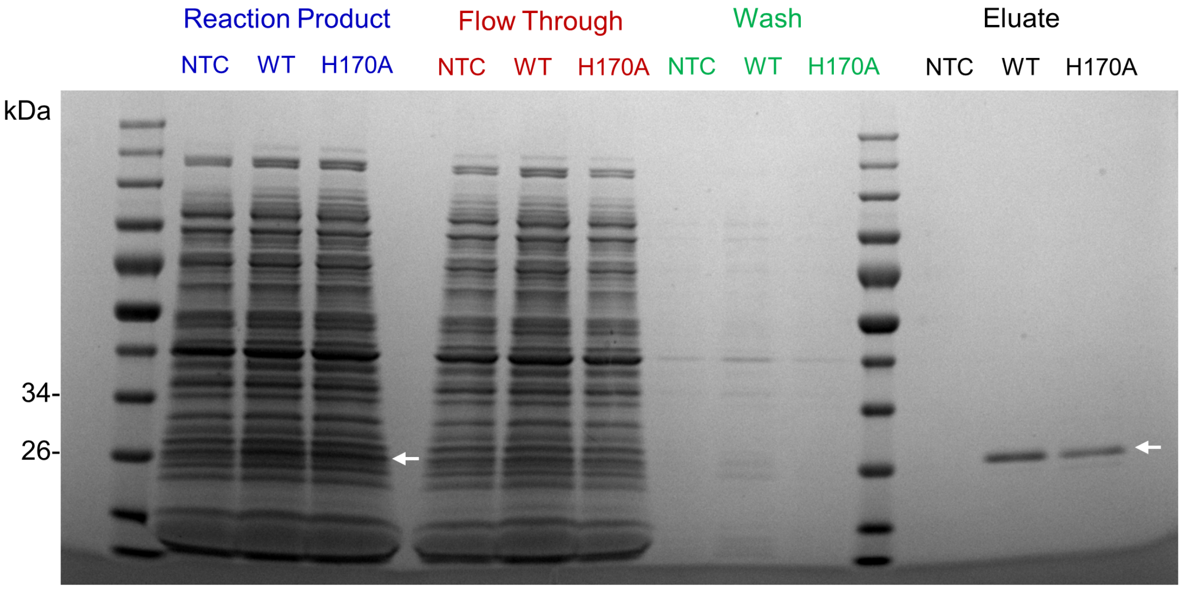


Figure S1: **SDS–PAGE analysis of cell-free expressed products and His-tag purified proteins.** Reaction products, flow-through, wash fractions, and eluted proteins were resolved by SDS–PAGE and stained with Coomassie blue. Lanes correspond to no-template control (NTC), wild-type HRP (WT), and H170A mutant, as indicated. The expected band for WT and H170A (∼32 kDa, arrows) is visible in the reaction products and can be clearly seen in the elution fractions after His-tag purification, confirming successful translation of both proteins in the cell-free system.

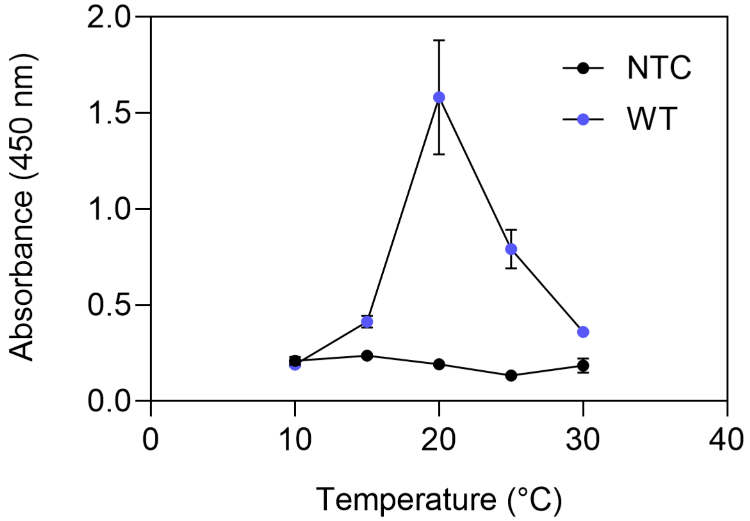


Figure S2: **Temperature optimisation of HRP expression in the cell-free system.** HRP activity was measured at different incubation temperatures by spectrophotometric detection of TMB oxidation. Wild-type HRP (WT, blue) shows maximal activity at 20 °C, whereas the no-template control (NTC, black) exhibits negligible signal across all tested temperatures.

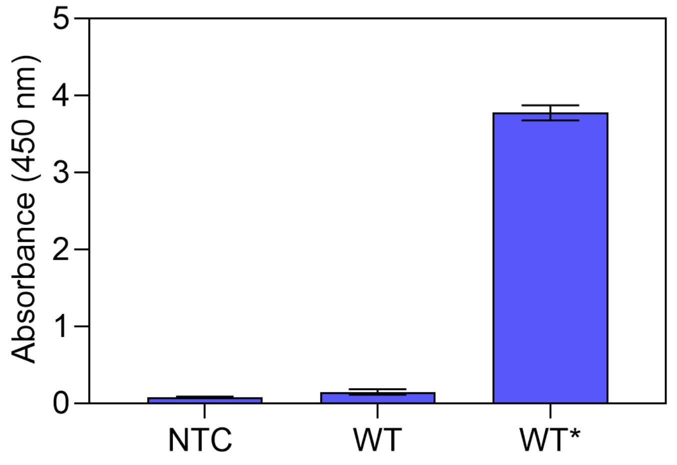


Figure S3: **Optimisation of cofactors for HRP expression in the cell-free system.** HRP was expressed at its optimal incubation temperature (20 °C) under different co-factor conditions and quantified by spectrophotometric detection of TMB oxidation at 450 nm following a 500-fold dilution of reaction products. WT indicates the conditions before optimisation (hemin : 10 $\mu$M, $Ca^{2+}$ : 2 mM, $Mg^{2+}$ : 5 mM), while WT* represents the final optimised condition (hemin : 15 $\mu$M, $Ca^{2+}$ : 0.5 mM, $Mg^{2+}$ : 2 mM). NTC shows a negligible background signal.

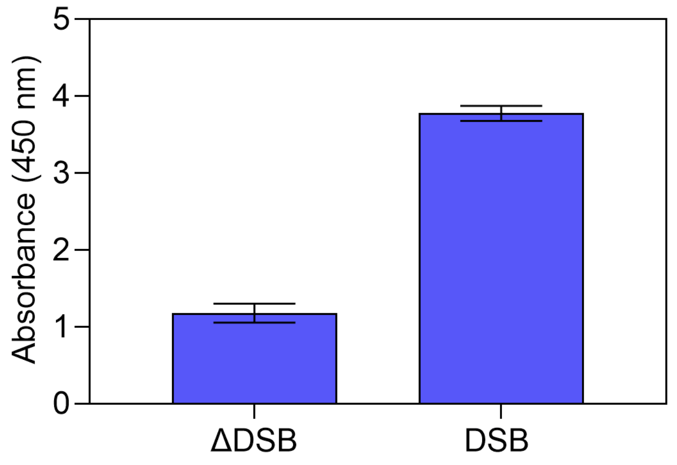


Figure S4: **Effect of disulfide bond enhancers (DSB) on HRP expression in the cell-free system.** HRP activity was measured under optimized conditions (20 °C with adjusted hemin, $Ca^{2+}$, and $Mg^{2+}$ concentrations) in the presence (DSB) or absence ($\Delta$DSB) of commerical disulfide bond enhancers. Activity was quantified by spectrophotometric detection of TMB oxidation following a 500-fold dilution of reaction products. Inclusion of DSB resulted in markedly higher HRP activity compared to $\Delta$DSB.

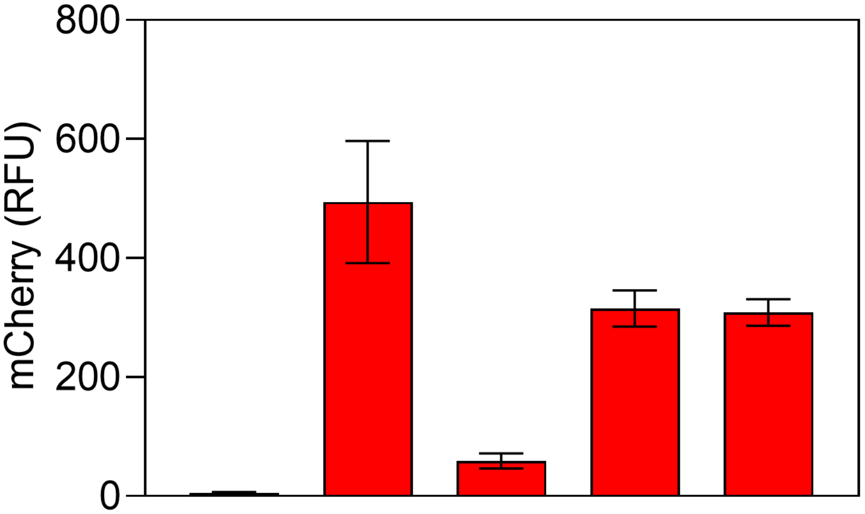


| | | | | | |
|---|---|---|---|---|---|
| T7-TetO-mCherry | - | + | + | + | + |
| TetR repressor | - | - | + | + | - |
| aTc activator | - | - | - | + | + |

Figure S5: **Optimization of a tetracycline-responsive genetic circuit using mCherry as reporter.** Fluorescence mCherry output was measured under different genetic circuit conditions to assess the requirement for reporter DNA, TetR repressor, and aTc activator. Reactions were performed with 10 nM T7–TetO mCherry DNA, 200 nM TetR protein, and 1 $\mu$M aTc in a total reaction volume of 10 $\mu$L, as indicated in the table below the graph.

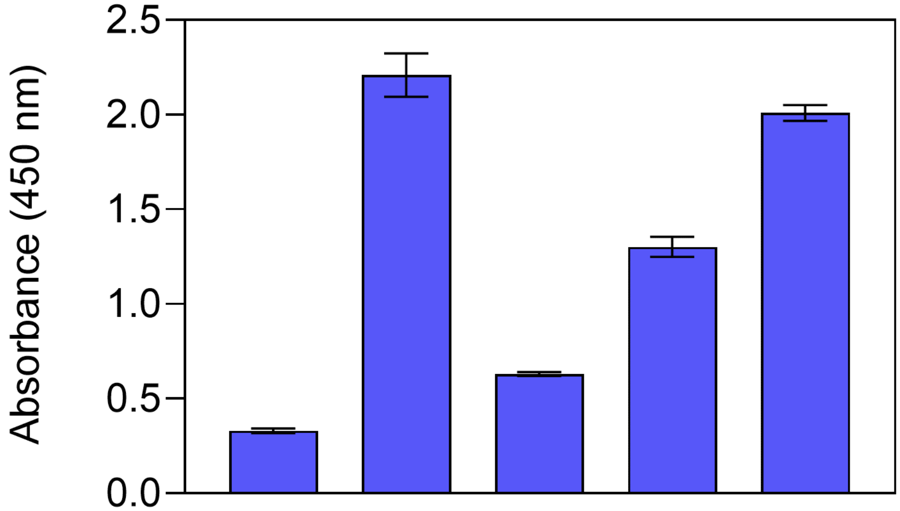


| | | | | | |
|---|---|---|---|---|---|
| T7-TetO-HRP | - | + | + | + | + |
| TetR repressor | - | - | + | + | - |
| aTc activator | - | - | - | + | + |

Figure S6: **Implementation of a tetracycline-responsive genetic circuit using HRP as reporter.** HRP activity was quantified under different genetic circuit conditions by spectrophotometric detection of TMB oxidation. Reactions contained 10 nM T7–TetO HRP DNA, 200 nM TetR protein, and 1 $\mu$M aTc activator in a total reaction volume of 10 $\mu$L, as indicated in the table below the graph.

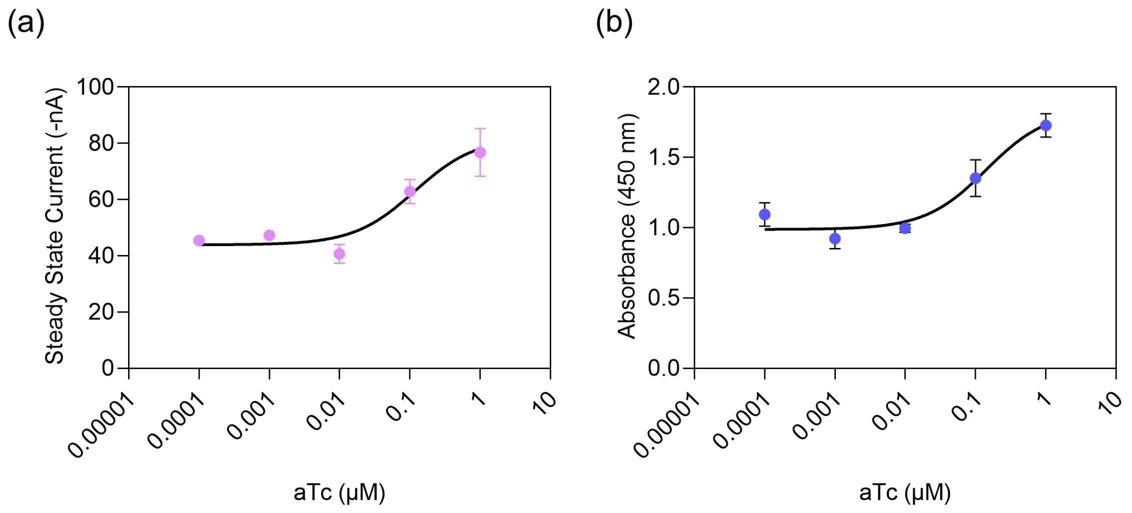


Figure S7: **Dose-response characterization of the aTc-inducible sensor using HRP as a reporter.** (a) Electrochemical response showing steady-state current as a function of increasing aTc concentration. (b) Colorimetric response measured as absorbance following TMB oxidation. Both datasets exhibit a sigmoidal response, indicating dose-dependent induction of HRP expression with increasing aTc concentrations.

### 2 Supplementary Tables

Table S1: Primers used for linear PCR amplification

| **Primer** | **Sequence (5'–3')** |
|---|---|
| **Forward** | TGCTGCAAGGCGATTAAGTT |
| **Reverse** | TGTAGGCATAGGCTTGGTTA |

Table S2: Composition of the cell-free reaction mixture.

| **Components** | **Final Concentration in Reaction** |
|---|---|
| Crude | 10 mg/mL |
| Energy Solution | See caption below |
| Amino Acid | See caption below |
| PEG 8000 | 2 mM |
| Magnesium Glutamate | variable |
| Potassium Glutamate | 100 mM |
| Maltose | 15 mM |
| DTT | 1.5 mM |
| Chi-6 | 5 $\mu$M |
| DNA Template | 10 nM |

The mastermix for all the TX-TL reactions was made in a 1.5 ml microcentrifuge tube. Additive components were thawed on ice before adding to the tube. The energy solution and amino acid solution were prepared and used as previously described [1]. In the reactions for HRP, magnesium glutamate was added at 2 mM and reactions were aliquoted into PCR tubes and incubated in the thermocycler (ProFlex PCR System), whereas for mCherry, magnesium glutamate was added at 8 mM, and reactions were incubated in the 96-well plate in the plate reader.
