## Supplementary Sequence Information for "Horseradish peroxidase as an electrochemical reporter protein for cell-free biosensors"

The DNA sequences used in this study are summarised below, with individual elements highlighted (Green is T7 promoter; Red is Tet operator; Blue is the insert; and Orange is 6xHis C-tag). Each construct includes 250 base pairs of flanking regions on either side of the insert, and the primers used for linear DNA preparation are also provided. A complete plasmid map is shown at the bottom.

| Template | Sequence (5’-3’) |
|---|---|
| T7_HRP | TGCTGCAAGGCGATTAAGTTGGGTAACGCCAGGGTTTTCCCAGTCACG<br>ACGTTGTAAAACGACGGCCAGTGCCAAGCTTGCATGCAAGGAGATGGC<br>GCCCAACAGTCCCCCGGCCACGGGGCCTGCCACCATACCCACGCCGA<br>AACAAGCGCTCATGAGCCCGAAGTGGCGAGCCCGATCTTCCCCATCG<br>GTGATGTCGGCGATATAGGCGCCAGCAACCGCACCTGTGGCGCCGGT<br>GATGCCGGCCACGATGCGTCCGGCGTAGAGGATCGAGATCTCGATCC<br>CGCGAAATTAATACGACTCACTATAGGGAGACCACAACGGTTTCCCTCT<br>AGAAATAATTTTGTTTAACTTTAAGAAGGAGATATACCATGCAACTCACC<br>CCAACTTTCTACGACAATTCATGCCCGAACGTTAGCAACATTGTCCGCG<br>ACACCATCGTAAACGAACTGCGTTCTGATCCGCGTATTGCTGCGTCCAT<br>CCTGCGCCTGCACTTCCACGATTGTTTCGTTAACGGATGCGACGCGTC<br>TATCCTGCTGGACAACACCACCTCCTTCCGTACCGAAAAGGATGCGTT<br>CGGCAACGCCAACTCCGCGCGGTTTCCCAGTTATCGACCGCATGAA<br>AGCCGCTGTAGAATCCGCCTGTCCGCGTACCGTATCTTGCGCAGACCT<br>CCTGACCATCGCGGCCCAGCAGAGCGTTACTCTAGCAGGTGGCCCGT<br>CTTGGCGTGTTCCGCTGGGTCGTCGTGATTCTCTACAGGCGTTCCTGG<br>ATCTGGCCAACGCAAATCTGCCAGCTCCGTTCTTCACCCTGCCGCAGC<br>TGAAAGATAGCTTCCGTAACGTTGGCCTGAACCGTTCATCCGATCTGGT<br>GGCGTTGTCTGGTGGTCACACCTTCGGGAAAAACCAGTGCCGTTTCAT<br>CATGGACCGCCTGTATAACTTCTCGAACACCGGTCTGCCGGACCCGAC<br>CCTGAACACCACCTATTTGCAGACTCTGCGTGGGCTGTGCCCGCTGAA<br>CGGTAACCTGTCCGCGCTGGTTGACTTCGATCTGCGTACTCCGACCAT<br>CTTCGATAACAAATACTACGTTAACCTGGAAGAACAGAAGGGCCTGATT<br>CAGTCTGACCAGGAGCTGTTCTCCTCCCCGAACGCGACCGACACCATC<br>CCGCTGGTTCGTAGCTTCGCGAACAGCACGCAGACTTTCTTCAACGCT<br>TTCGTAGAGGCTATGGACCGTATGGGTAACATTACCCCGCTGACCGGT<br>ACGCAGGGACAGATCCGCCTGAACTGCCGCGTGGTTAACTCCAACTCC<br>CACCACCACCACCATCACGTGTAAGATCCGGCTGCTAACAAAGCCCGA<br>AAGGAAGCTGAGTTGGCTGCTGCCACCGCTGAGCAATAACTAGCATAA<br>CCCCTTGGGGCCTCTAAACGGGTCTTGAGGGGTTTTTTGCTGAAAGGA<br>GGAACTATATCCGGATATCCACAGGACGGGTGTGGTCGCCATGATCGC<br>GTAGTCGATAGTGGCTCCAAGTAGCGAAGCGAGCAGGACTGGGCGGC<br>GGCCAAAGCGGTCGGACAGTGCTCCGAGAACGGGTGCGCATAGAAAT<br>TGCATCAACGCATATAGCGCTAGCAGCACGCCATAGTGACTGGCGATG<br>CTGTCGGAATGGACGATATCCCGCAAGAGGCCCGGCAGTACCGGCAT<br>AACCAAGCCTATGCCTACA |
| T7_H170A | TGCTGCAAGGCGATTAAGTTGGGTAACGCCAGGGTTTTCCCAGTCACG<br>ACGTTGTAAAACGACGGCCAGTGCCAAGCTTGCATGCAAGGAGATGGC<br>GCCCAACAGTCCCCCGGCCACGGGGCCTGCCACCATACCCACGCCGA<br>AACAAGCGCTCATGAGCCCGAAGTGGCGAGCCCGATCTTCCCCATCG<br>GTGATGTCGGCGATATAGGCGCCAGCAACCGCACCTGTGGCGCCGGT |

| | |
|---|---|
| | GATGCCGGCCACGATGCGTCCGGCGTAGAGGATCGAGATCTCGATCC<br>CGCGAAATTAATACGACTCACTATAGGGAGACCACAACGGTTTCCCTCT<br>AGAAATAATTTTGTTTAACTTTAAGAAGGAGATATACCATGCAACTCACC<br>CCAACTTTCTACGACAATTCATGCCCGAACGTTAGCAACATTGTCCGCG<br>ACACCATCGTAAACGAACTGCGTTCTGATCCGCGTATTGCTGCGTCCAT<br>CCTGCGCCTGCACTTCCACGATTGTTTCGTTAACGGATGCGACGCGTC<br>TATCCTGCTGGACAACACCACCTCCTTCCGTACCGAAAAGGATGCGTT<br>CGGCAACGCCAACTCCGCGCGGTTTCCCAGTTATCGACCGCATGAA<br>AGCCGCTGTAGAATCCGCCTGTCCGCGTACCGTATCTTGCGCAGACCT<br>CCTGACCATCGCGGCCCAGCAGAGCGTTACTCTAGCAGGTGGCCCGT<br>CTTGGCGTGTTCCGCTGGGTCGTCGTGATTCTCTACAGGCGTTCCTGG<br>ATCTGGCCAACGCAAATCTGCCAGCTCCGTTCTTCACCCTGCCGCAGC<br>TGAAAGATAGCTTCCGTAACGTTGGCCTGAACCGTTCATCCGATCTGGT<br>GGCGTTGTCTGGTGGTGCAACCTTCGGGAAAAACCAGTGCCGTTTCAT<br>CATGGACCGCCTGTATAACTTCTCGAACACCGGTCTGCCGGACCCGAC<br>CCTGAACACCACCTATTTGCAGACTCTGCGTGGGCTGTGCCCGCTGAA<br>CGGTAACCTGTCCGCGCTGGTTGACTTCGATCTGCGTACTCCGACCAT<br>CTTCGATAACAAATACTACGTTAACCTGGAAGAACAGAAGGGCCTGATT<br>CAGTCTGACCAGGAGCTGTTCTCCTCCCCGAACGCGACCGACACCATC<br>CCGCTGGTTCGTAGCTTCGCGAACAGCACGCAGACTTTCTTCAACGCT<br>TTCGTAGAGGCTATGGACCGTATGGGTAACATTACCCCGCTGACCGGT<br>ACGCAGGGACAGATCCGCCTGAACTGCCGCGTGGTTAACTCCAACTCC<br>CACCACCACCACCATCACGTGTAAGATCCGGCTGCTAACAAAGCCCGA<br>AAGGAAGCTGAGTTGGCTGCTGCCACCGCTGAGCAATAACTAGCATAA<br>CCCCTTGGGGCCTCTAAACGGGTCTTGAGGGGTTTTTTGCTGAAAGGA<br>GGAACTATATCCGGATATCCACAGGACGGGTGTGGTCGCCATGATCGC<br>GTAGTCGATAGTGGCTCCAAGTAGCGAAGCGAGCAGGACTGGGCGGC<br>GGCCAAAGCGGTCGGACAGTGCTCCGAGAACGGGTGCGCATAGAAAT<br>TGCATCAACGCATATAGCGCTAGCAGCACGCCATAGTGACTGGCGATG<br>CTGTCGGAATGGACGATATCCCGCAAGAGGCCCGGCAGTACCGGCAT<br>AACCAAGCCTATGCCTACA |
| T7_tetO_HRP | GAAAGGGGGATGTGCTGCAAGGCGATTAAGTTGGGTAACGCCAGGGT<br>TTTCCCAGTCACGACGTTGTAAAACGACGGCCAGTGCCAAGCTTGCAT<br>GCAAGGAGATGGCGCCCAACAGTCCCCCGGCCACGGGGCCTGCCACC<br>ATACCCACGCCGAAACAAGCGCTCATGAGCCCGAAGTGGCGAGCCCG<br>ATCTTCCCCATCGGTGATGTCGGCGATATAGGCGCCAGCAACCGCACC<br>TGTGGCGCCGGTGATGCCGGCCACGATGCGTCCGGCGTAGAGGATCG<br>AGATCTCGATCCCGCGAAATTAATACGACTCACTATAGGGTCCCTATCA<br>GTGATAGAGACCGCTCTAGAAATAATTTGTTTAACTTTAAGAAGGAGAT<br>ATACATATGCAACTCACCCCAACTTTCTACGACAATTCATGCCCGAACG<br>TTAGCAACATTGTCCGCGACACCATCGTAAACGAACTGCGTTCTGATCC<br>GCGTATTGCTGCGTCCATCCTGCGCCTGCACTTCCACGATTGTTTCGTT<br>AACGGATGCGACGCGTCTATCCTGCTGGACAACACCACCTCCTTCCGT<br>ACCGAAAAGGATGCGTTCGGCAACGCCAACTCCGCGCGGTTTCCC<br>AGTTATCGACCGCATGAAAGCCGCTGTAGAATCCGCCTGTCCGCGTAC<br>CGTATCTTGCGCAGACCTCCTGACCATCGCGGCCCAGCAGAGCGTTAC<br>TCTAGCAGGTGGCCCGTCTTGGCGTGTTCCGCTGGGTCGTCGTGATTC<br>TCTACAGGCGTTCCTGGATCTGGCCAACGCAAATCTGCCAGCTCCGTT<br>CTTCACCCTGCCGCAGCTGAAAGATAGCTTCCGTAACGTTGGCCTGAA<br>CCGTTCATCCGATCTGGTGGCGTTGTCTGGTGGTCACACCTTCGGGAA |

| | |
|---|---|
| | AAACCAGTGCCGTTTCATCATGGACCGCCTGTATAACTTCTCGAACACC<br>GGTCTGCCGGACCCGACCCTGAACACCACCTATTTGCAGACTCTGCGT<br>GGGCTGTGCCCGCTGAACGGTAACCTGTCCGCGCTGGTTGACTTCGAT<br>CTGCGTACTCCGACCATCTTCGATAACAAATACTACGTTAACCTGGAAG<br>AACAGAAGGGCCTGATTCAGTCTGACCAGGAGCTGTTCTCCTCCCCGA<br>ACGCGACCGACACCATCCCGCTGGTTCGTAGCTTCGCGAACAGCACG<br>CAGACTTTCTTCAACGCTTTCGTAGAGGCTATGGACCGTATGGGTAACA<br>TTACCCCGCTGACCGGTACGCAGGGACAGATCCGCCTGAACTGCCGC<br>GTGGTTAACTCCAACTCCCACCACCACCACCATCACGTGTAAGATCCG<br>GCTGCTAACAAAGCCCGAAAGGAAGCTGAGTTGGCTGCTGCCACCGCT<br>GAGCAATAACTAGCATAACCCCTTGGGGCCTCTAAACGGGTCTTGAGG<br>GGTTTTTTGCTGAAAGGAGGAACTATATCCGGATATCCACAGGACGGG<br>TGTGGTCGCCATGATCGCGTAGTCGATAGTGGCTCCAAGTAGCGAAGC<br>GAGCAGGACTGGGCGGCGGCCAAAGCGGTCGGACAGTGCTCCGAGA<br>ACGGGTGCGCATAGAAATTGCATCAACGCATATAGCGCTAGCAGCACG<br>CCATAGTGACTGGCGATGCTGTCGGAATGGACGATATCCCGCAAGAGG<br>CCCGGCAGTACCGGCATAACCAAGCCTATGCCTACAGCATCCAGGGTG<br>AC |
| T7_tetO_mCherry | GAAAGGGGGATGTGCTGCAAGGCGATTAAGTTGGGTAACGCCAGGGT<br>TTTCCCAGTCACGACGTTGTAAAACGACGGCCAGTGCCAAGCTTGCAT<br>GCAAGGAGATGGCGCCCAACAGTCCCCCGGCCACGGGGCCTGCCACC<br>ATACCCACGCCGAAACAAGCGCTCATGAGCCCGAAGTGGCGAGCCCG<br>ATCTTCCCCATCGGTGATGTCGGCGATATAGGCGCCAGCAACCGCACC<br>TGTGGCGCCGGTGATGCCGGCCACGATGCGTCCGGCGTAGAGGATCG<br>AGATCTCGATCCCGCGAAATTAATACGACTCACTATAGGGTCCCTATCA<br>GTGATAGAGACCGCTCTAGAAATAATTTTGTTTAACTTTAAGAAGGAGAT<br>ATACATATGGTGAGCAAGGGCGAGGAGGATAACATGGCCATCATCAAG<br>GAGTTCATGCGCTTCAAGGTGCACATGGAGGGCTCCGTGAACGGCCA<br>CGAGTTCGAGATCGAGGGCGAGGGCGAGGGCCGCCCCTACGAGGGC<br>ACCCAGACCGCCAAGCTGAAGGTGACCAAGGGTGGCCCCCTGCCCTT<br>CGCCTGGGACATCCTGTCCCCTCAGTTCATGTACGGCTCCAAGGCCTA<br>CGTGAAGCACCCCGCCGACATCCCCGACTACTTGAAGCTGTCCTTCCC<br>CGAGGGCTTCAAGTGGGAGCGCGTGATGAACTTCGAGGACGGCGGCG<br>TGGTGACCGTGACCCAGGACTCCTCCCTGCAGGACGGCGAGTTCATCT<br>ACAAGGTGAAGCTGCGCGGCACCAACTTCCCCTCCGACGGCCCCGTA<br>ATGCAGAAGAAGACCATGGGCTGGGAGGCCTCCTCCGAGCGGATGTA<br>CCCCGAGGACGGCGCCCTGAAGGGCGAGATCAAGCAGAGGCTGAAGC<br>TGAAGGACGGCGGCCACTACGACGCTGAGGTCAAGACCACCTACAAG<br>GCCAAGAAGCCCGTGCAGCTGCCCGGCGCCTACAACGTCAACATCAA<br>GTTGGACATCACCTCCCACAACGAGGACTACACCATCGTGGAACAGTA<br>CGAACGCGCCGAGGGCCGCCACTCCACCGGCGGCATGGACGAGCTG<br>TACAAGCACCACCACCACCACCATCACGTGTAAGATCCGGCTGCTAACAAA<br>GCCCGAAAGGAAGCTGAGTTGGCTGCTGCCACCGCTGAGCAATAACTA<br>GCATAACCCCTTGGGGCCTCTAAACGGGTCTTGAGGGGTTTTTTGCTG<br>AAAGGAGGAACTATATCCGGATATCCACAGGACGGGTGTGGTCGCCAT<br>GATCGCGTAGTCGATAGTGGCTCCAAGTAGCGAAGCGAGCAGGACTG<br>GGCGGCGGCCAAAGCGGTCGGACAGTGCTCCGAGAACGGGTGCGCA<br>TAGAAATTGCATCAACGCATATAGCGCTAGCAGCACGCCATAGTGACT<br>GGCGATGCTGTCGGAATGGACGATATCCCGCAAGAGGCCCGGCAGTA<br>CCGGCATAACCAAGCCTATGCCTACAGCATCCAGGGTGAC |

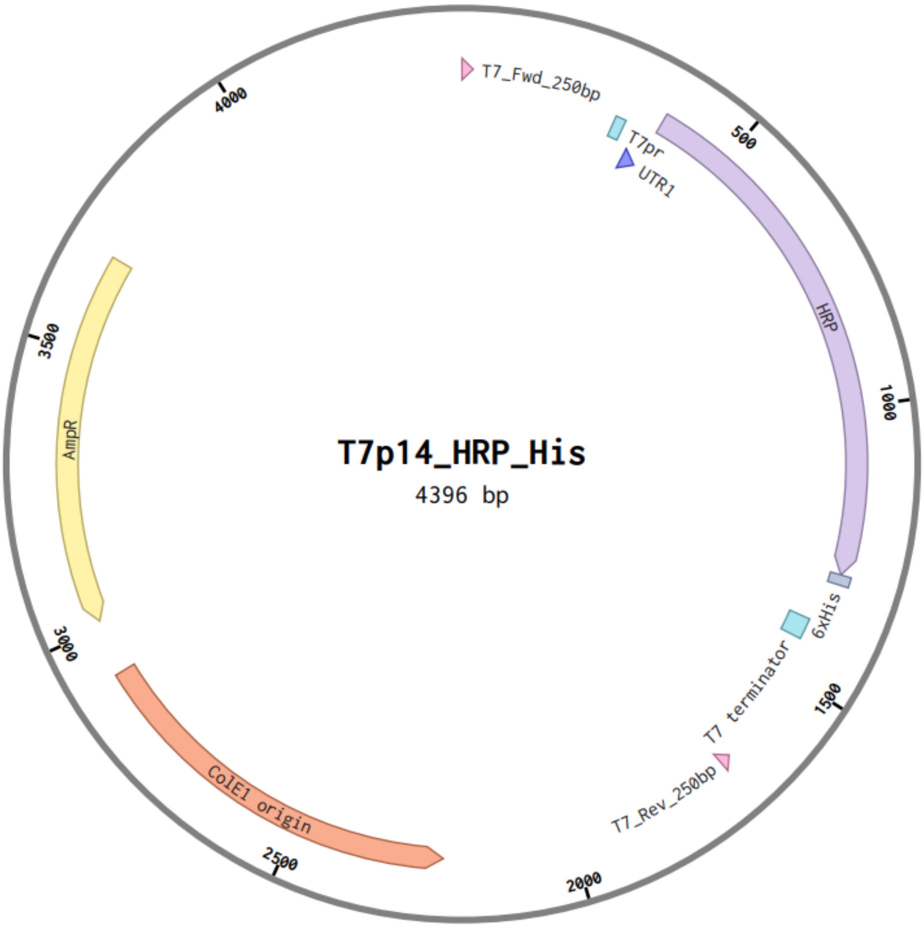

T7p14_HRP_His
4396 bp
T7_Fwd_250bp
T7pr
UTR1
HRP
6xHis
T7 terminator
T7_Rev_250bp
ColE1 origin
AmpR
500
1000
1500
2000
2500
3000
3500
4000
